# Generation of porcine PK-15 cells lacking the *Ifnar1* gene to optimize the efficiency of viral isolation

**DOI:** 10.1101/2023.07.28.550964

**Authors:** Maya Shofa, Akatsuki Saito

**Affiliations:** Department of Veterinary Science, Faculty of Agriculture, University of Miyazaki, Miyazaki, Miyazaki 8892192, Japan; Graduate School of Medicine and Veterinary Medicine, University of Miyazaki, Miyazaki, Miyazaki 8891692, Japan; Center for Animal Disease Control, University of Miyazaki, Miyazaki, Miyazaki 8892192, Japan

## Abstract

Because pigs are intermediate or amplifying hosts for several zoonotic viruses, the pig-derived PK-15 cell line is an indispensable tool for studying viral pathogenicity and developing treatments, vaccines, and preventive measures to mitigate the risk of disease outbreaks. However, we must consider the possibility of contamination by type I interferons (IFNs), such as IFNα and IFNβ, or IFN-inducing substances, such as virus-derived double-stranded RNA or bacterial lipopolysaccharides, in clinical samples, leading to lower rates of viral isolation. In this study, we aimed to generate a PK-15 cell line that can be used to isolate viruses from clinical samples carrying a risk of contamination by IFN-inducing substances. To this end, we depleted the IFN alpha and beta receptor subunit 1 (*Ifnar1*) gene in PK-15 cells using a clustered regularly interspaced short palindromic repeats (CRISPR)/CRISPR-associated protein 9 method. Treatment of PK-15 cells lacking *Ifnar1* with IFNβ resulted in no inhibitory effects on viral infection by a lentiviral vector, influenza virus, and Akabane virus. These results demonstrate that PK-15 cells lacking *Ifnar1* could represent a valuable and promising tool for viral isolation, vaccine production, and virological investigations.

## Introduction

Type I interferons (IFNs) such IFNα and IFNβ are significant components of innate immunity against viral infections [1,2]. IFNs exert their biological effects by binding to IFN alpha and beta receptor subunit 1 (IFNAR1) on the cellular surface and subsequently activating Janus kinase 1 (JAK1) and tyrosine kinase 2 [3]. Signal transducer and activator of transcription (STAT) transduction mediates cytokine responses. The JAK–STAT signaling pathway triggers the intracellular IFN signaling cascade, which eventually induces the expression of IFN-stimulated genes (ISGs) to generate an antiviral state [3,4].

Despite the importance of the IFN cascade as the first line of defense against viral infections, induction of this cascade can perturb the isolation of infectious viruses from clinical samples if there is contamination by IFN-inducing agents such as virus-derived double-stranded RNA (dsRNA) or bacterial lipopolysaccharide. These IFN-inducing agents are recognized by sensor proteins in host cells such as melanoma differentiation-associated protein 5 [5], retinoic acid-inducible gene-I [6,7], and Toll-like receptors [8–11], leading to IFN production, and the subsequent induction of an antiviral state. In this aim, Vero cells have been widely used to isolate viruses from clinical samples [12]. One major reason to use Vero cells for viral isolation is that these cells cannot produce IFNs [13,14]. However, because Vero cells were derived from an African green monkey (*Cercopithecus aethiops*), these cells might not be optimal for the isolation and replication of viruses from other species.

Pigs are intermediate or amplifying hosts of several zoonotic viruses such as influenza virus, Japanese encephalitis virus, Nipah virus, and coronavirus [15]. Furthermore, pigs are essential animals globally. Thus, preventing viral diseases in pigs is critical for sustainable agriculture. Several of life-threading viral diseases affect pigs, including African swine fever virus, classical swine fever virus [16], porcine circovirus type 2 [17], porcine transmissible gastroenteritis virus [18], porcine parvovirus [19], foot-and-mouth disease virus [20], and pseudorabies virus [21]. Therefore, suitable cell lines derived from pigs are critical to isolate these viruses from clinical samples. The porcine kidney-derived cell line PK-15 has been used for this purpose and for vaccine development. Unlike Vero cells, which lack IFN production [14,22], PK-15 cells have an intact pathway induced by IFNs. Therefore, if the clinical samples contain IFNs or IFN-inducing substances, such as virus-derived dsRNA, or bacterial lipopolysaccharide, PK-15 cells produce IFNs upon stimulation, leading to the induction of ISGs and unsuccessful virus isolation and replication.

In this study, we sought to knock out *Ifnar1* in PK-15 cells to overcome this limitation. We used a clustered regularly interspaced short palindromic repeats (CRISPR)/CRISPR-associated protein 9 (Cas9) method to knock out *Ifnar1* in PK-15 cells. We compared the replication of influenza A virus (IAV) and Akabane virus (AKAV) between normal and *Ifnar1* knockout (k/o) PK-15 cells. PK-15 cells lacking *Ifnar1* exhibited robust viral replication in the presence of pig IFNβ. Together, *Ifnar1* k/o PK-15 cells could represent a valuable tool for viral isolation, thereby optimizing vaccine production, and virological investigation.

## Materials and Methods

### Plasmids

The psPAX2-IN/HiBiT and pWPI-Luc2 plasmids were kind gifts from Dr. Kenzo Tokunaga [23]. pMD2.G was a gift from Dr. Didier Trono (Cat# 12259; http://n2t.net/addgene:12259; RRID: Addgene_12259), and pSpCas9(BB)-2A-Puro (PX459) V2.0 was a gift from Dr. Feng Zhang (Addgene, Watertown, MA, USA, plasmid #62988; http://n2t.net/addgene:62988; RRID: Addgene_62988). pGP plasmid and pDON-5 Neo-ZsGreen plasmid were described previously [24].

To generate a retroviral vector expressing pig IFNAR1, the coding sequence of pig IFNAR1 was synthesized according to the amino acid sequences deposited in GenBank (Acc. Num. NM_213772.1) with codon optimization to pig cells (Integrated DNA Technologies, Inc., Coralville, IA, USA). The synthesized DNA sequence is summarized in S1 File. We next cloned the synthesized DNA into the pDON-5 Neo-vector (TaKaRa, Kusatsu, Japan, Cat# 3657), which was pre-linearized with *Not*I-HF (New England Biolabs [NEB], Ipswich, MA, USA, Cat# R3189L) and *Bam*HI-HF (NEB, Cat# R3136L) using an In-Fusion HD Cloning Kit (TaKaRa, Cat# Z9633N). The plasmid was amplified using NEB 5-alpha F′ Iq Competent *E. coli* (High Efficiency) (NEB, Cat# C2992H) and extracted using the PureYield Plasmid Miniprep System (Promega, Madison, WI, USA, Cat# A1222). The sequence of the plasmid was verified by sequencing using a SupreDye v3.1 Cycle Sequencing Kit (M&S TechnoSystems, Osaka, Japan, Cat# 063001) with a Spectrum Compact CE System (Promega). To generate a pDON-5 Neo-vector expressing IFNAR1 with the W70C mutation, we performed mutagenesis by overlapping PCR using PrimeSTAR GXL DNA Polymerase (TaKaRa, Cat# R050A) together with the primers IFNAR-F (5′-CGTGGGCCCGCGGCCGCGCCACCATG-3’) and W70C-R (5′-GAGCTTGATgCAGTTATCCATTCCAGTGA-3′) to amplify the 5′ fragment and the primers W70C-F (5′-ATGGATAACTGcATCAAGCTCCCTGGATG-3′) and IFNAR1-R (5′-TAACGTCGACGGATCCCTACACTGC-3′) to amplify the 3′ fragment. These fragments were mixed and amplified with the primers IFNAR1-F and IFNAR1-R. The resultant IFNAR1 fragment encoding the W70C mutation was cloned into the pDON-5 Neo-vector as previously described.

### Cell culture

Human kidney-derived Lenti-X 293T cells (TaKaRa, Cat# Z2180N) and porcine kidney-derived PK-15 cells (Japanese Collection of Research Bioresources Cell Bank, Cat# JCRB9040) were maintained in Dulbecco’s modified Eagle medium (DMEM; Nacalai Tesque, Kyoto, Japan, Cat# 08458-16) supplemented with 10% fetal bovine serum (FBS) and 1× penicillin–streptomycin (Pe/St; Nacalai Tesque, Cat# 09367-34).

### Viruses

IAV (H1N1) strain A/PR/8/34 (American Type Culture Collection, Manassas, VA, USA, Cat# VR-95) was used in this study. IAV was propagated in specific pathogen-free chicken embryonated eggs. AKAV (TS-C2 vaccine strain) was purchased from Kyoto Biken Laboratories (Kyoto, Japan). AKAV was propagated in HmLu-1 cells maintained in DMEM supplemented with 2% FBS and 1% Pe/St. All experiments were performed in a biosafety level 2 facility.

### Rescue of HIV-1–based reporter virus

To rescue an HIV-1–based lentiviral vector expressing the luciferase 2 reporter gene, Lenti-X 293T cells were co-transfected with the psPAX2-IN/HiBiT, pWPI-Luc2, and pMD2.G plasmids using TransIT-293 Transfection Reagent (TaKaRa, Cat# V2700) in Opti-MEM I Reduced Serum Medium (Thermo Fisher Scientific, Waltham, MA, USA, Cat# 31985062). The supernatant was filtered 2 days after transfection.

### Design of single guide RNAs (sgRNAs)

Specific sgRNAs targeting pig *Ifnar1* and *Stat2* were designed using the online CRISPR Design Tool (https://crispr.dbcls.jp). The sequence of sgRNAs used in this study is summarized in Table 1. Two partially complementary oligos were used to generate the sgRNA scaffold. The complementary oligos were mixed and heated at 95°C for 5 min, followed by incubation at room temperature for 1 h for oligo annealing. The annealed oligos were diluted 250-fold with water and used for ligation with the pSpCas9(BB)-2A-Puro (PX459) V2.0 vector (Addgene, Cat# 62988), which was predigested with *Bbs*I-HF (NEB, Cat# R3539L) using a DNA Ligation Kit (Mighty Mix, TaKaRa, Cat# 6023). The ligated constructs were then transformed in NEB 5-alpha F′ Iq Competent *E. coli* (High Efficiency). The plasmids were analyzed using a primer (5′-ACTATCATATGCTTACCGTAAC-3′) to verify the sequence.

**Table 1.**
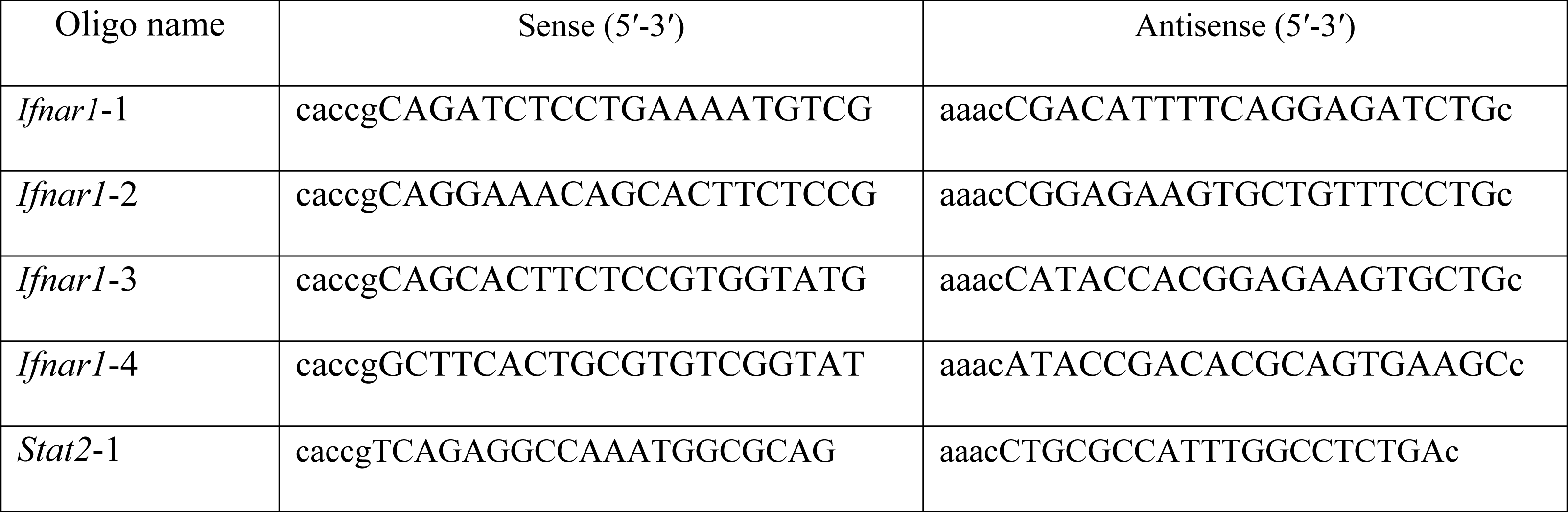
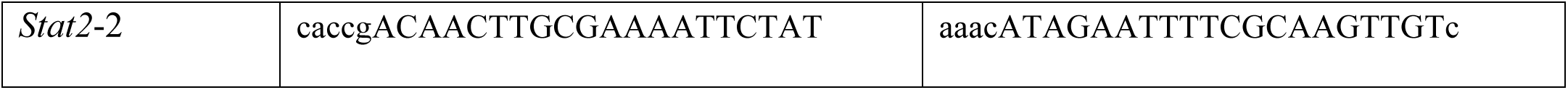
The oligonucleotides used to generate the sgRNA scaffolds.

### Generation of PK-15 cells lacking *Ifnar1* or *Stat2*

To generate k/o cells, PK-15 cells were transfected with PX459-*Ifnar1*-k/o or PX459-*Stat2*-k/o plasmids using the TransIT-X2 Dynamic Delivery System (TaKaRa, Cat# V6100) according to the manufacturer’s instructions. At 48 h after transfection, cells were treated with 5 μg/mL puromycin (InvivoGen, San Diego, CA, USA, Cat# ant-pr-1). After 1 week of incubation, single-cell clones were sorted in a 96-well plate using a Cell Sorter SH800S (Sony Biotechnology, Inc., San Diego, CA, USA). The single-cell clones were characterized by infection with an HIV-based lentiviral vector.

### Infection of PK-15 cells lacking *Ifnar1* or *Stat2* with lentiviral vectors for screening

Normal PK-15 cells, PK-15 *Ifnar1* k/o cells, or PK-15 *Stat2* k/o cells were plated (1 × 10^4^ cells/well) in a 96-well plate and treated with 0, 10, or 100 ng/mL pig IFNβ (Kingfisher Biotech, St. Paul, MN, USA, Cat# RP0011S-025). After overnight incubation, cells were infected with an HIV-1–based reporter virus expressing luciferase 2. Two days after infection, cells were lysed using the Britelite plus reporter gene assay system (PerkinElmer, Waltham, MA, USA, Cat# 6066769), and the luminescent signal was measured using a GloMax Explorer Multimode Microplate Reader (Promega).

### Measurement of ISG expression

Normal PK-15 cells, PK-15 *Ifnar1* k/o cells, or PK-15 *Stat2* k/o cells were plated in a 96-well plate in sextuplicate as previously described and treated with 0 or 100 ng/mL IFNβ. After overnight incubation, total RNA was collected using a CellAmp Direct RNA Prep Kit for RT-PCR (Real Time) (TaKaRa, Cat# 3732) according to the manufacturer’s instructions. Porcine myxovirus resistance 1 (*Mx1*), *ISG15*, and *Viperin* messenger RNA (mRNA) levels were measured by a qRT-PCR assay using the One Step TB Green PrimeScript PLUS RT-PCR Kit (Perfect Real Time) (TaKaRa, Cat# RR096A). The PCR protocol was 42°C for 5 min, 95 °C for 10 s, and 40 cycles of 95°C for 5 s and 60°C for 34 s. Porcine *Mx1*, *ISG15*, and *Viperin* mRNA levels were normalized to those of porcine β-actin, which was used as an endogenous control (ΔΔCt method). The specific sequence primers for ISG mRNA levels were described previously [4]. The primer sets are listed in Table 2.

**Table 2.**
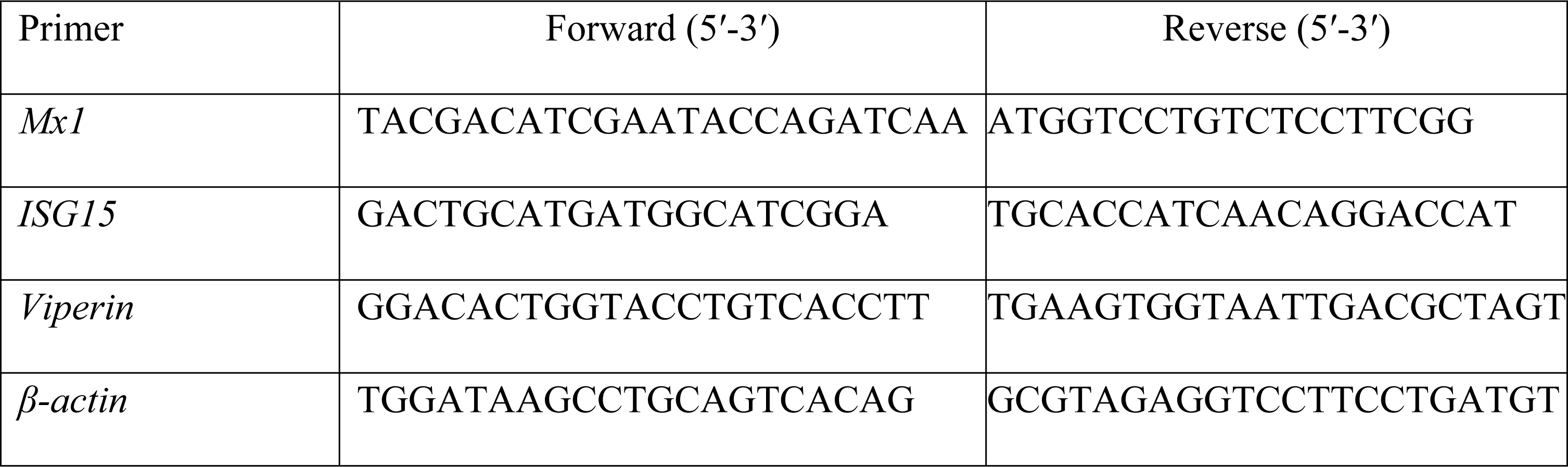
The oligonucleotides used to quantify ISG mRNA levels.

### Western blotting

To evaluate ISG15 expression, the pelleted cells were lysed in 2× Bolt LDS sample buffer (Thermo Fisher Scientific, Cat# B0008) containing 2% β-mercaptoethanol (Bio-Rad, Hercules, CA, USA, Cat# 1610710) and incubated at 70°C for 10 min. ISG15 expression was evaluated using SimpleWestern Abby (ProteinSimple, San Jose, CA, USA) with anti-ISG15 rabbit polyclonal antibody (Cell Signaling Biotechnology, Danvers, MA, USA, Cat# 2743S, ×250) and an Anti-Rabbit Detection Module (ProteinSimple, Cat# DM-001). The amount of input protein was visualized using a Total Protein Detection Module (ProteinSimple, Cat# DM-TP01).

### Virus replication assay

Normal PK-15 cells, PK-15 *Ifnar1* k/o cells, or PK-15 *Stat2* k/o cells were plated at 1 × 10^4^ cells/well in a 96-well plate and treated with 0 or 100 ng/mL pig IFNβ. After overnight culture, cells were infected with IAV or AKAV. After 2 h of incubation, cells were washed, and fresh DMEM supplemented with 2% FBS and 1× Pe/St was added to each well with or without 100 ng/mL pig IFNβ. For experiments using IAV, cells were maintained in DMEM supplemented with 10% FBS, 1× Pe/St, and 1 μg/mL tosylsulfonyl phenylalanyl chloromethyl ketone-treated trypsin (Sigma-Aldrich, St. Louis, MO, USA, Cat# 4352157) [25,26]. The supernatant was harvested 2 days after infection and mixed with 2× RNA lysis buffer (2% Triton X-100, 50 mM KCl, 100 mM Tris-HCl [pH 7.4], 40% glycerol, 0.4 U/μL Recombinant RNase Inhibitor [TaKaRa, Cat# 2313A]) as described previously [27]. The replication levels of IAV and AKAV were determined by qRT-PCR using primer pairs (Table 3) as described previously [28]. The analysis was performed in a QuantStudio 5 Real-Time PCR System (Applied Biosystems, Waltham, MA, USA) using a One Step TB Green PrimeScript PLUS RT-PCR Kit (Perfect Real Time). The relative levels of viral RNA were calculated using the ΔΔCt method.

**Table 3.**
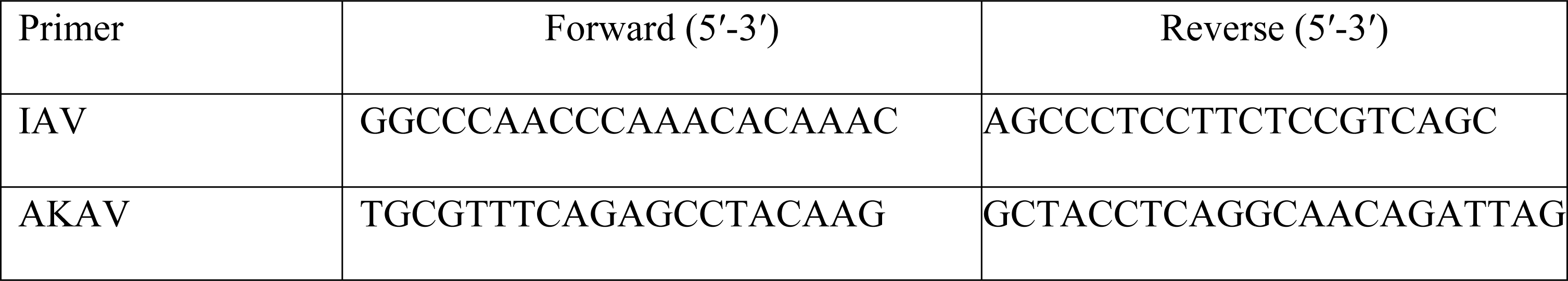
The oligonucleotides used for the virus replication assay.

### Reconstitution of IFNAR1 molecules in PK-15 cells lacking *Ifnar1* or *Stat2*

To rescue retroviral vectors expressing pig wild-type (WT) IFNAR1, pig IFNAR1 carrying the W70C mutation, or ZsGreen protein, 3 × 10^6^ Lenti-X 293T cells were co-transfected with the pGP vector, pDON-5 Neo-IFNAR1 (WT), pDON-5 Neo-IFNAR1 (W70C), or pDON-5 Neo-ZsGreen, and pMD2.G plasmids using TransIT-293 Transfection Reagent in Opti-MEM I Reduced Serum Medium. The supernatant was filtered 2 days after transfection. To avoid contamination by residual plasmid DNA, viral stocks were treated with 4 U/mL TURBO DNase (Thermo Fisher Scientific, Cat# AM2238) at 37°C for 60 min.

To reconstitute IFNAR1, PK-15 cells lacking *Ifnar1*, or *Stat2* were infected with retroviral vectors expressing pig IFNAR1 (WT), pig IFNAR1 (W70C), or ZsGreen protein. After 48 h of incubation, the cells were treated with 1 mg/mL G418 disulfate aqueous solution (Nacalai Tesque, Cat# 09380-86) to deplete non-transduced cells. Cells were maintained until cells transduced with ZsGreen-expressing virus became >90% ZsGreen-positive. PK-15 *Ifnar1* k/o or PK-15 *Stat2* k/o cells infected with retroviral vectors expressing IFNAR1 (WT), IFNAR1 (W70C), or ZsGreen protein were plated at 1 × 10^4^ cells/well. After 24 h, total RNA was extracted using a CellAmp Direct RNA Prep Kit for RT-PCR (Real Time) to measure the *Ifnar1* expression. We used a forward primer (5′-GCTGAGGACAAGGCGATTAT-3′) and a reverse primer (5′-GGAGTACACGAATGAGGATGAG-3′) based on the codon-optimized sequence of pig *Ifnar1* (S1 File). Expression was measured as previously described. To assess antiviral activity in reconstituted cells, cells were treated with 0 or 100 ng/mL IFNβ for 24 h, and the viral infection experiment was performed as previously described. The mRNA expression of IGSs in reconstituted cells was determined as previously described.

### Statistical analysis

Differences between treated and untreated cells were evaluated by an unpaired, two-tailed Student’s *t*-test. Multiple comparisons were evaluated by one-way ANOVA followed by Tukey’s test. *p* ≤ 0.05 indicated statistical significance. The test was performed using GraphPad Prism 9 software v9.1.1 (GraphPad, San Diego, CA, USA).

## Results

### Generation and screening of PK-15 cells lacking *Ifnar1*

To generate PK-15 cells lacking *Ifnar1*, we designed sgRNAs targeting pig *Ifnar1* using pig genome information deposited in GenBank (NM_213772.1). Furthermore, we designed sgRNAs targeting pig *Stat2* because STAT2 is a key molecule involved in IFN-mediated antiviral responses [29–31]. PK-15 cells were transfected with PX459 plasmids harboring these sgRNAs, and the single-cell colony was isolated using a limiting dilution method. Single clones of PK-15 *Ifnar1* k/o or PK-15 *Stat2* k/o cells were tested via infection by HIV-1–based reporter virus after treating cells with pig IFNβ. Normal PK-15 cells efficiently blocked infection by the HIV-1–based reporter virus (Fig 1). By contrast, PK-15 *Ifnar1* and PK-15 *Stat2* k/o cells did not resist HIV-1–based reporter virus infection (Fig 1), suggesting that cells lacking *Ifnar1* or *Stat2* are not responsive to IFNβ treatment.

**Fig 1.**
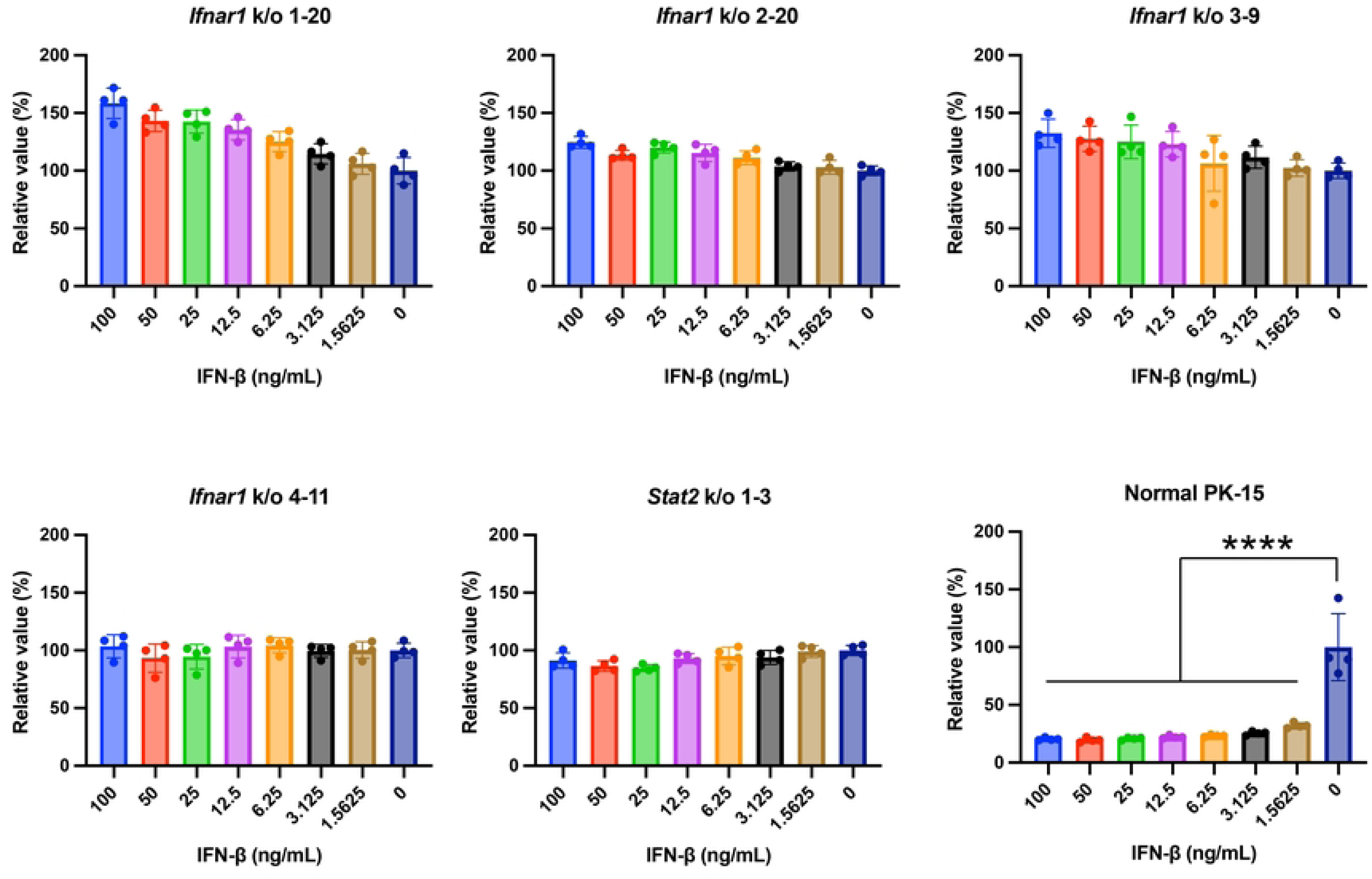
Generation and screening of PK-15 cells lacking *Ifnar1*. Normal PK-15 cells, PK-15 *Ifnar1* k/o cells, or PK-15 *Stat2* k/o cells treated with various concentrations of pig IFNβ were infected with HIV-1–based reporter virus. Infectivity was calculated as relative light units (RLU) 2 days after infection. Relative infectivity was calculated in comparison that that in untreated cells. The results are presented as the mean and standard deviation of quadruplicate measurements from one assay, and they are representative of at least three independent experiments. Differences were examined by one-way ANOVA followed by Tukey’s test. \*\*\*\**p* < 0.0001.

### PK-15 cells lacking *Ifnar1* lost the ability to induce ISGs upon IFNβ treatment

Activation of the type I IFN signaling pathway triggers a complex cascade of events that culminate in the expression of a broad array of ISGs, contributing to an antiviral state in infected cells. To characterize PK-15 *Ifnar1* k/o and PK-15 *Stat2* k/o cells, we evaluated the mRNA expression of three ISGs (*Mx1*, *ISG15*, and *Viperin*) upon IFNβ treatment. Treatment of normal PK-15 cells with 100 ng/mL pig IFNβ robustly induced ISG expression (over 100-fold) compared to that in untreated cells (Fig 2a and 2b). Conversely, both PK-15 *Ifnar1* k/o and PK-15 *Stat2* k/o cells displayed no or marginal induction of these ISGs upon IFNβ treatment (Fig 2a and 2b). Furthermore, we tested the induction of ISG15 protein upon IFNβ treatment by western blotting. In normal PK-15 cells, we observed a significant induction of ISG15 upon IFNβ treatment (Fig 2c), whereas ISG15 expression was not induced by IFNβ treatment in PK-15 *Ifnar1* k/o and PK-15 *Stat2* k/o cells (Fig 2c). These results demonstrated that PK-15 cells lacking *Ifnar1* or *Stat2* genes were unresponsive to IFNβ treatment.

**Fig 2.**
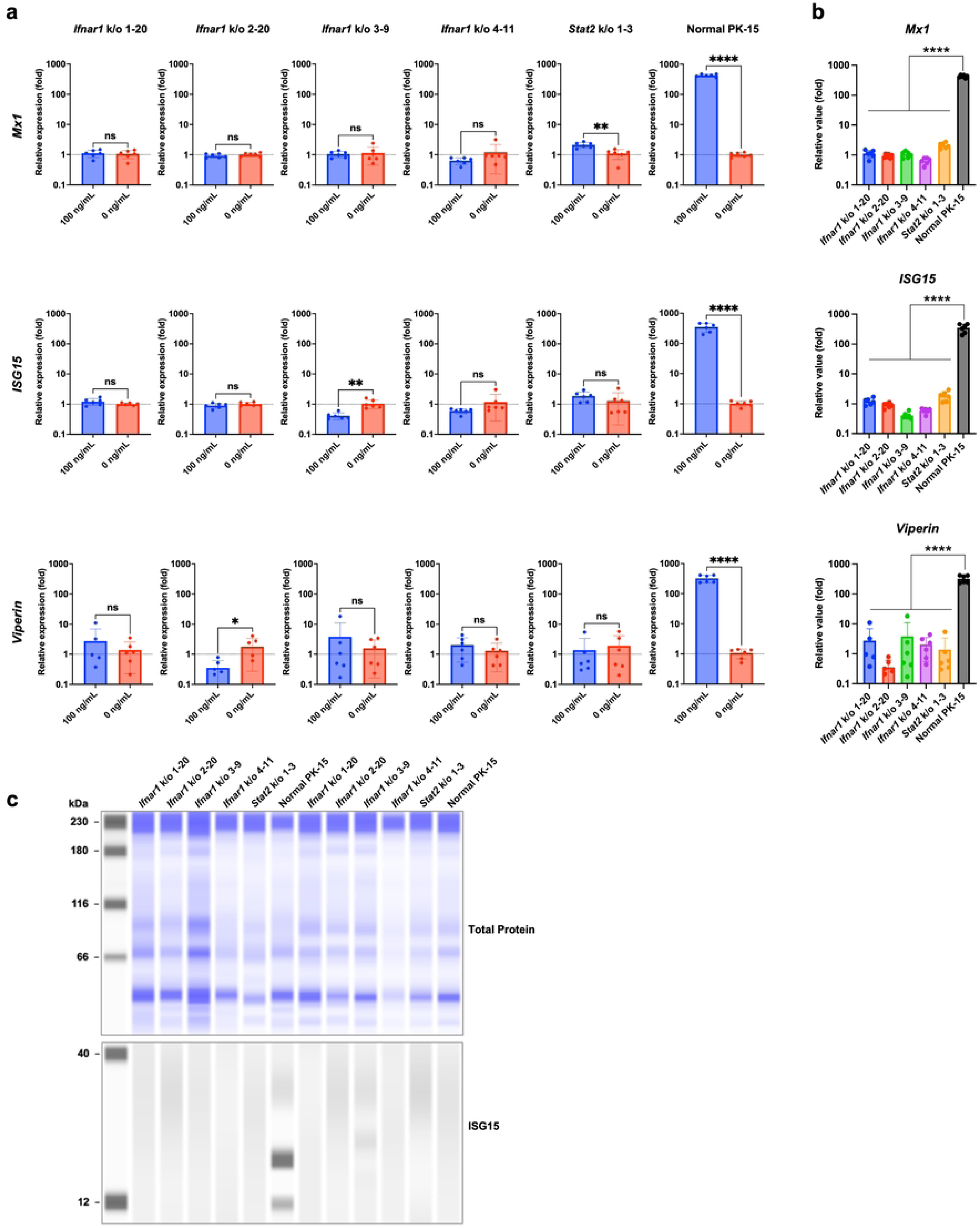
PK-15 cells lacking *Ifnar1* exhibited a loss of ISG induction upon IFNβ treatment. **(a)** Induction of ISG mRNAs in normal PK-15 cells, PK-15 *Ifnar1* k/o cells, and PK-15 *Stat2* k/o cells was measured by qRT-PCR 1 day after IFNβ treatment. The results are presented as the mean and standard deviation of sextuplicate measurements from one assay, and they are representative of at least three independent experiments. Differences between 100 ng/mL IFNβ-treated cells and untreated cells were examined by a two-tailed, unpaired Student’s *t*-test. \*\*\*\**p* < 0.0001, ns (not significant). **(b)** The data presented in Fig 2a were used to compare ISG induction among normal PK-15 cells, PK-15 *Ifnar1* k/o cells, and PK-15 *Stat2* k/o cells. Multiple comparisons were examined by one-way ANOVA followed by Tukey’s test. \*\*\*\**p* < 0.0001. **(c)** Expression of ISG15 protein in normal PK-15 cells, PK-15 *Ifnar1* k/o cells, and PK-15 *Stat2* k/o cells was determined by western blotting.

### PK-15 cells lacking *Ifnar1* are susceptible to viral infection in the presence of IFNβ

Next, we examined whether *Ifnar1* k/o augments the susceptibility of PK-15 cells to viral infection in the presence of IFNβ. We used PK-15 *Ifnar1* k/o (Clone #4-11) and PK-15 *Stat2* k/o cells (Clone #1-2) for further analyses. IAV replication was significantly inhibited in normal PK-15 cells treated with IFNβ (Fig 3a). By contrast, IAV replication was not inhibited in PK-15 *Ifnar1* k/o and PK-15 *Stat2* k/o cells upon IFNβ treatment (Fig 3a). Previous research demonstrated the inter-species transmission of AKAV from cows to pigs [32]. Therefore, the prevalence, and pathogenicity of AKAV in the pig population should be investigated. To test the usefulness of PK-15 *Ifnar1* k/o cells for this purpose, we tested the replication of AKAV in PK-15 cells treated with pig IFNβ (Fig 3b). Whereas AKAV replication was blocked in normal PK-15 cells treated with IFNβ, there was no inhibition of AKAV replication in PK-15 *Ifnar1* k/o and PK-15 *Stat2* k/o cells treated with IFNβ (Fig 3b). These results demonstrated that both PK-15 *Ifnar1* k/o and PK-15 *Stat2* k/o cells supported viral replication in the presence of IFNβ.

**Fig 3.**
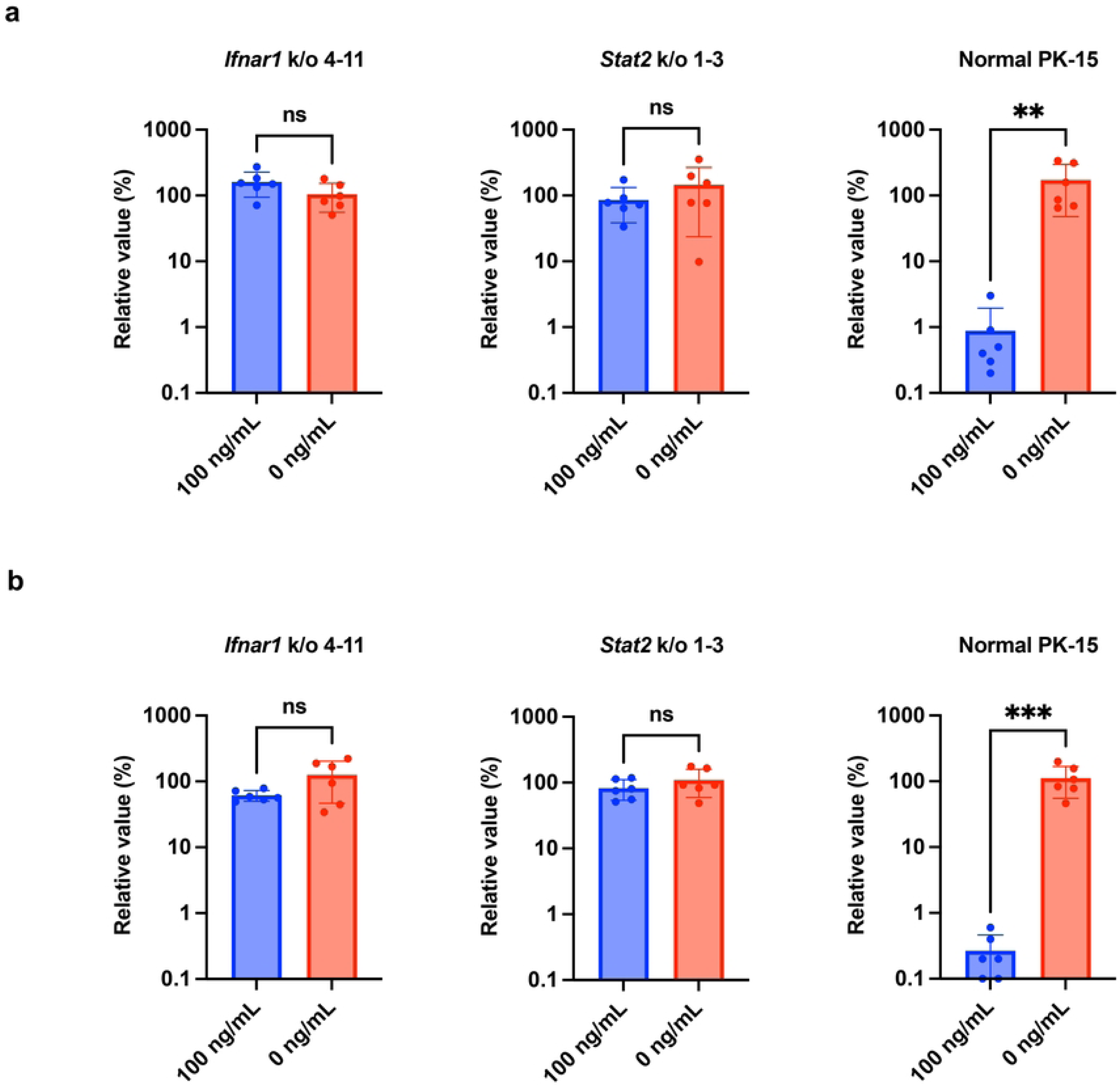
PK-15 cells lacking *Ifnar1* are susceptible to viral infection in the presence of IFNβ. Replication of IAV (**a**) and AKAV (**b**) was examined in normal PK-15 cells, PK-15 *Ifnar1* k/o cells, and PK-15 *Stat2* k/o cells. Cells were pretreated with 0 or 100 ng/mL IFNβ before infection. Viral RNA levels in the culture supernatant were quantified by qRT-PCR 2 days after infection. The relative values were calculated in comparison to that in non-pretreated cells. The results are presented as the mean and standard deviation of sextuplicate measurements from one assay. Differences between 100 ng/mL IFNβ-treated cells and untreated cells were examined by a two-tailed, unpaired Student’s *t-*test. \*\*\**p* < 0.001, \*\**p* < 0.05, ns (not significant).

### Reconstitution of *Ifnar1* in PK-15 *Ifnar1* k/o cells restored the response to IFNβ

Off-target effects represent an important concern of the CRISPR/Cas9 method [33]. To elucidate the specificity of *Ifnar1* k/o, we reconstituted IFNAR1 by infecting PK-15 *Ifnar1* k/o and PK-15 *Stat2* k/o cells with retroviral vectors expressing pig IFNAR1. In this experiment, we used both IFNAR1 (WT) and mutant IFNAR1 carrying the W70C mutation (W73C in human IFNAR1), IFNAR1 (W70C), which was previously reported to be associated with impaired activity of human IFNAR1 [34]. Although we attempted to detect reconstituted IFNAR1 by western blotting, we failed to detect pig IFNAR1 using three commercial anti-human IFNAR1 antibodies. Therefore, we used qRT-PCR for detection. *Ifnar1* k/o cells reconstituted with IFNAR1 (WT) or IFNAR1 (W70C) displayed significantly lower Ct values than control cells (Fig 4a), suggesting that IFNAR1 molecules were successfully reconstituted in these cells. We performed an infection assay in the reconstituted cells using AKAV. The result demonstrated that PK-15 *Ifnar1* k/o cells reconstituted with IFNAR1 (WT) resisted viral infection similarly as normal PK-15 cells (Fig 4b). Conversely, PK-15 *Ifnar1* k/o cells reconstituted with IFNAR1 (W70C) did not resist viral infection, suggesting that the restoration of antiviral activity required reconstitution with IFNAR1 (WT). As expected, neither IFNAR1 (WT) nor IFNAR1 (W70C) restored the resistance of PK-15 *Stat2* k/o cells to viral infection (Fig 4b). Supporting the result of the infection activity, ISG expression was induced by IFNβ treatment in PK-15 *Ifnar1* k/o cells reconstituted with IFNAR1 (WT) but not in the other cells (Fig 4b). These results suggested that the PK-15 *Ifnar1* k/o cells generated in this study exhibit a specific deletion of *Ifnar1*.

**Fig 4.**
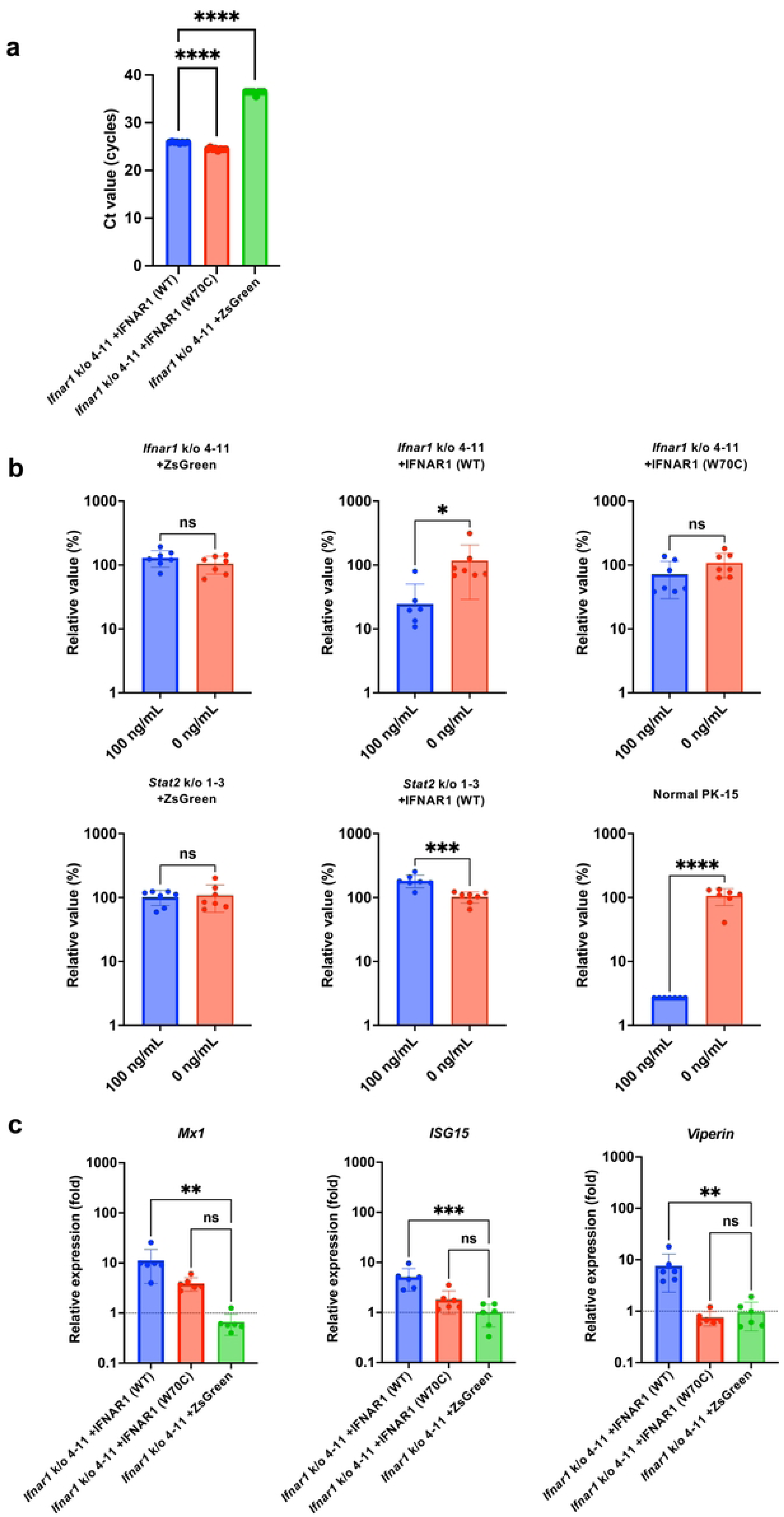
Reconstitution of IFNAR1 in PK-15 Ifnar1 k/o cells restored the response to IFNβ. **(a)** PK-15 *Ifnar1* k/o cells were infected with retroviral vectors expressing IFNAR1 (WT), IFNAR1 (W70C), or ZsGreen protein. After selection with G418, *Ifnar1* expression was measured by qRT-PCR 1 day after infection. The results are presented as the mean and standard deviation of octuplicate measurements from one assay, and they are representative of at least three independent experiments. Multiple comparisons were examined by one-way ANOVA followed by Tukey’s test. \*\*\*\**p* < 0.0001. **(b)** PK-15 *Ifnar1* k/o and PK-15 *Stat2* k/o cells infected with retroviral vectors expressing IFNAR1 (WT), IFNAR1 (W70C), or ZsGreen protein were treated with 0 or 100 ng/mL IFNβ for 24 h. Cells were superinfected with AKAV. Viral RNA levels in the culture supernatant were quantified by qRT-PCR 2 days after infection. The relative values were calculated in comparison to that in non-pretreated cells. The results are presented as the mean and standard deviation of septuplicate measurements from one assay, and they are representative of at least three independent experiments. Differences between 0 and 100 ng/mL IFNβ-treated cells were examined by a two-tailed, unpaired Student’s *t-*test. \*\*\*\**p* < 0.0001, \*\*\**p* < 0.001, \**p* < 0.05, ns (not significant). **(c)** Induction of ISG mRNAs in PK-15 *Ifnar1* k/o cells infected with retroviral vectors expressing IFNAR1 (WT), IFNAR1 (W70C), or ZsGreen protein was measured by qRT-PCR 1 day after IFNβ treatment. The results are presented as the mean and standard deviation of sextuplicate measurements from one assay, and they are representative of at least three independent experiments. Multiple comparisons were examined by one-way ANOVA followed by Tukey’s test. \*\*\**p* < 0.001, \*\**p* < 0.01, ns (not significant).

## Discussion

In this study, we demonstrated that PK-15 *Ifnar1* k/o cells represent a promising tool for viral isolation from clinical samples, vaccine development, and virological investigation. Our results illustrated that PK-15 *Ifnar1* k/o cells were susceptible to HIV-1–based reporter virus infection upon IFNβ treatment. We observed marginal ISG expression upon IFNβ treatment in PK-15 *Ifnar1* k/o cells compared to that in normal PK-15 cells. This result was confirmed by viral replication assays using IAV and AKAV.

IFNs play major role in innate immunity. IFNs are essential molecules for inducing an antiviral state, thereby protecting virus-exposed cells, and neighboring cells. IFNs bind to IFNAR1 on the cell surface, initiating a downstream cascade of immune responses. Stimulation of this cascade leads to the induction of ISGs such as *Mx1*, *ISG15*, and *Viperin* [35]. These ISGs block several stages of viral replication including viral entry, viral genome replication, and progeny virion release [36–38]. Mx protein has antiviral effects against influenza virus [2,39,40] and bunyavirus [38,41]. In this study, robust ISG induction and inhibition of viral replication were observed in normal PK-15 cells treated with IFNβ, whereas these effects were cancelled by *Ifnar1* or *Stat2* k/o.

We examined the potential risk of off-target effects of gene editing in PK-15 *Ifnar1* k/o cells. Reconstitution of PK-15 *Ifnar1* k/o cells with WT IFNAR1, but not IFNAR1 carrying the W70C mutation, restored the resistance of the cells to viral infection and ISG induction. Our results are consistent with a previous report demonstrating that W70C mutation in IFNAR1 (W73C in humans) increased the severity of COVID-19 because of the loss of IFN-mediated immune responses [34].

In conclusion, our study demonstrated that PK-15 *Ifnar1* k/o cells can be used for viral isolation from samples with possible contamination by IFNs or IFN-inducing substances. We believe that PK-15 *Ifnar1* k/o cells will contribute to the isolation of several zoonotic diseases such as influenza virus, Japanese encephalitis virus, and Nipah virus. Furthermore, considering the importance of maintaining the health of pigs for sustainable agriculture, this cell line can be used to understand the prevalence and pathogenicity of life-threatening viral diseases in pigs, including African swine fever, and foot-and-mouth disease.

## Acknowledgments

pMD2.G was a gift from Dr. Didier Trono. pSpCas9(BB)-2A-Puro (PX459) V2.0 was a gift from Dr. Feng Zhang. psPAX2-IN/HiBiT and pWPI-Luc2 were kind gifts from Dr. Kenzo Tokunaga. We thank Dr. Hirohisa Mekata, Ms. Tomoko Nishiuchi, and Ms. Yuki Shibatani for their support. We additionally thank Enago (www.enago.com) for the English language review.

## Author information contributions

M.S. and A.S. designed experiments, performed experiments, analyzed results, and wrote the manuscript. All authors read and approved the manuscript.

## Supporting information

**S1 File.**
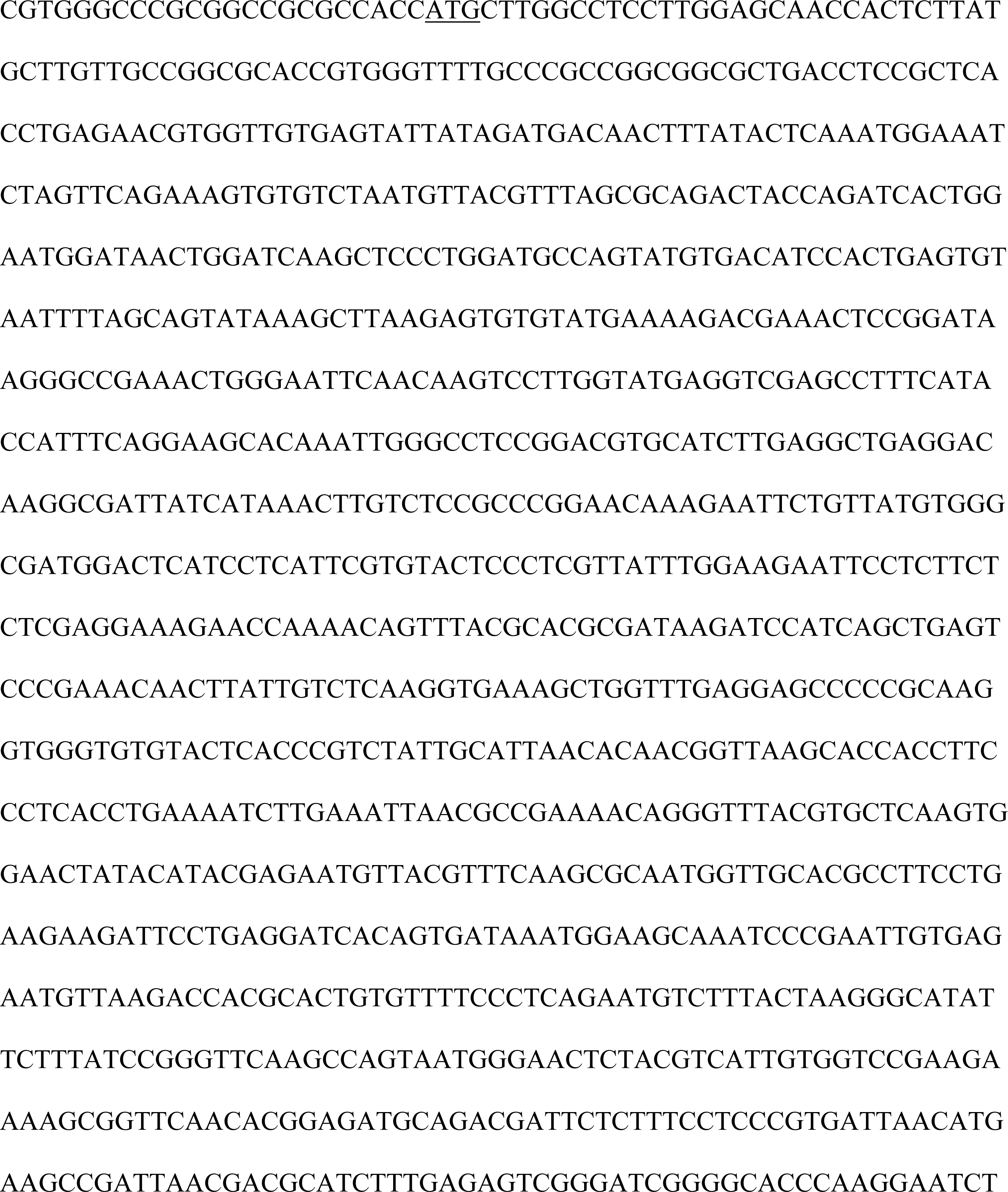
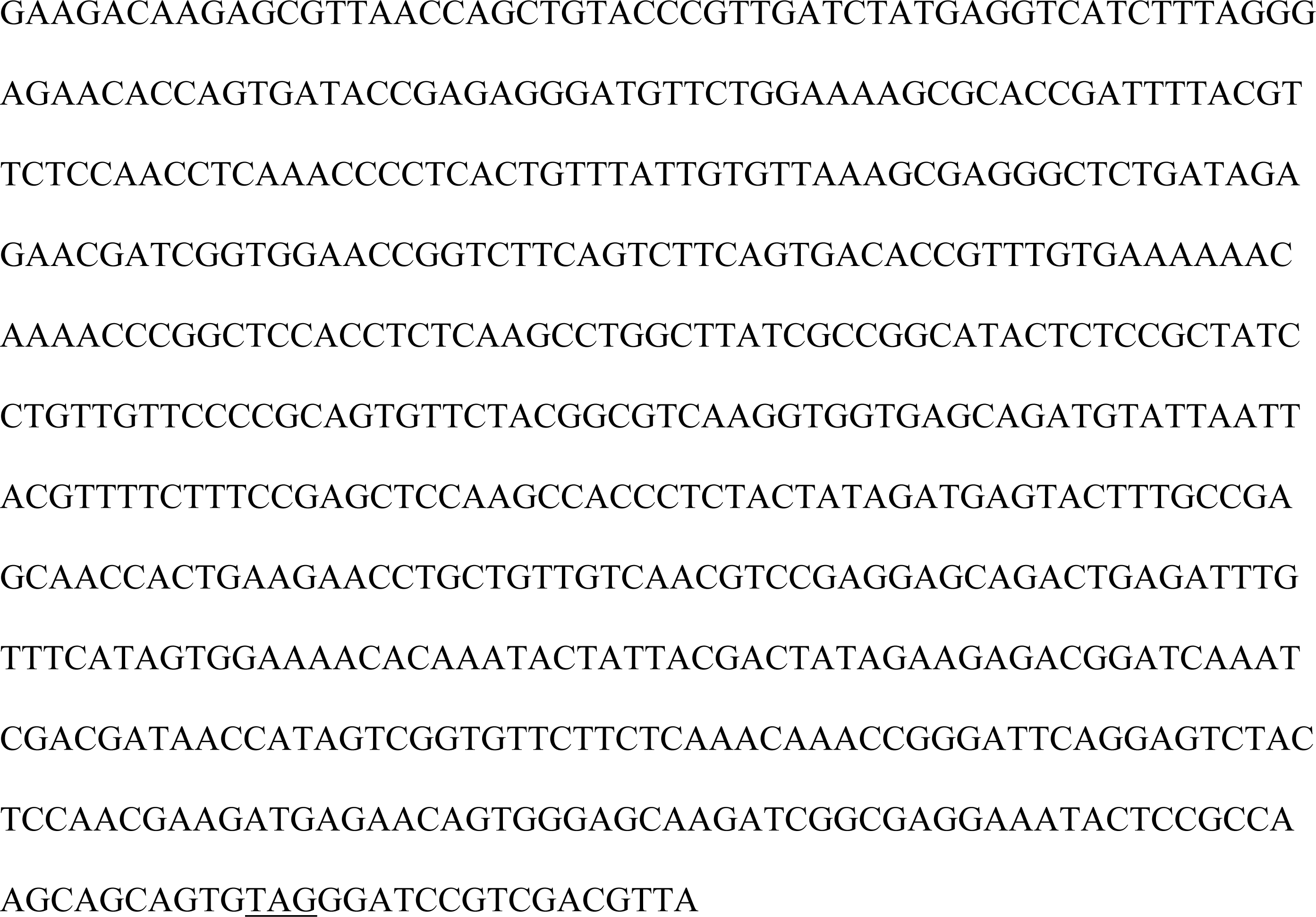
Synthesized DNA for generating plasmids expressing pig IFNAR1. The coding sequence of pig IFNAR1 was synthesized according to the amino acid sequence deposited in GenBank (Acc. Num. NM_213772.1) with codon optimization to pig cells. The start and stop codons are underlined.

